# Probing the biophysical constraints of SARS-CoV-2 spike N-terminal domain using deep mutational scanning

**DOI:** 10.1101/2022.06.20.496903

**Authors:** Wenhao O. Ouyang, Timothy J.C. Tan, Ruipeng Lei, Ge Song, Collin Kieffer, Raiees Andrabi, Kenneth A. Matreyek, Nicholas C. Wu

**Affiliations:** Department of Biochemistry, University of Illinois at Urbana-Champaign, Urbana, IL 61801, USA; Center for Biophysics and Quantitative Biology, University of Illinois at Urbana-Champaign, Urbana, IL 61801, USA; Department of Immunology and Microbiology, The Scripps Research Institute, La Jolla, CA 92037, USA; IAVI Neutralizing Antibody Center, The Scripps Research Institute, La Jolla, CA 92037, USA; Consortium for HIV/AIDS Vaccine Development (CHAVD), The Scripps Research Institute, La Jolla, CA 92037, USA; Department of Microbiology, University of Illinois at Urbana-Champaign, Urbana, IL 61801, USA; Department of Pathology, Case Western Reserve University School of Medicine, Cleveland, OH 44106, USA; Carl R. Woese Institute for Genomic Biology, University of Illinois at Urbana-Champaign, Urbana, IL 61801, USA; Carle Illinois College of Medicine, University of Illinois at Urbana-Champaign, Urbana, IL 61801, USA

## Abstract

Increasing the expression level of the SARS-CoV-2 spike (S) protein has been critical for COVID-19 vaccine development. While previous efforts largely focused on engineering the receptor-binding domain (RBD) and the S2 subunit, the N-terminal domain (NTD) has been long overlooked due to the limited understanding of its biophysical constraints. In this study, the effects of thousands of NTD single mutations on S protein expression were quantified by deep mutational scanning. Our results revealed that in terms of S protein expression, the mutational tolerability of NTD residues was inversely correlated with their proximity to the RBD and S2. We also identified NTD mutations at the interdomain interface that increased S protein expression without altering its antigenicity. Overall, this study not only advances the understanding of the biophysical constraints of the NTD, but also provides invaluable insights into S-based immunogen design.

## INTRODUCTION

The emergence of severe acute respiratory syndrome coronavirus-2 (SARS-CoV-2) has led to the coronavirus disease 2019 (COVID-19) pandemic^1, 2^. As the major antigen of SARS-CoV-2, spike (S) glycoprotein plays a critical role in facilitating virus entry^3, 4^. Therefore, antibodies to SARS-CoV-2 S are often neutralizing^5, 6^. SARS-CoV-2 S protein consists of an N-terminal S1 subunit, which is responsible for engaging the host receptor angiotensin-converting enzyme 2 (ACE2) via the receptor-binding domain (RBD), as well as a C-terminal S2 subunit, which mediates virus-host membrane fusion^4, 7, 8^. The S1 subunit also contains an N-terminal domain (NTD) in addition to the RBD^4, 7^. While the RBD is generally considered to be immunodominant, the NTD is also a target of neutralizing antibodies^9–11^. Structural studies revealed the presence of an antigenic supersite on the NTD that is frequently mutated in SARS-CoV-2 variants of concern (VOCs)^12–17^. In fact, amino acid mutations and indels rapidly accumulate within the NTD during the evolution of SARS-CoV-2 in human, at least partly due to the immune selection pressure^18^. On the other hand, antibodies to NTD epitopes that are conserved across VOCs have also been identified^16, 19^. Despite the importance of NTD in immune response against SARS-CoV-2, the biophysical constraints of NTD remain largely elusive.

COVID-19 vaccines, including both recombinant protein-based and mRNA-based, are proven to be highly protective against SARS-CoV-2 infection^20–23^. There is an inverse relationship between the production yield and cost of recombinant protein-based COVID-19 vaccines, such as that from Novavax, which showed promising results in phase 3 clinical trials^22^, as well as others that are in earlier phases of clinical trials^24^. High protein expression level is also believed to be critical for the effectiveness of mRNA vaccines^25^. As a result, identifying mutations that increase S protein expression are crucial for optimizing COVID-19 vaccines. While most studies focused on mutating the S2 subunit as well as the RBD to increase S protein expression^7, 26–29^, little effort has been spent on NTD due to the lack of understanding of its biophysical properties.

Phenotypes of numerous mutations can be measured in a massively parallel manner using deep mutational scanning, which combines saturation mutagenesis and next-generation sequencing^30^. Previous studies have applied deep mutational scanning to evaluate the effects of RBD mutations on protein expression, ACE2-binding affinity, and antibody escape^31–36^. Although deep mutational scanning of the RBD provided important insights into immunogen design and SARS-CoV-2 evolution^29, 31, 32, 35, 36^, similar studies on other regions of the S protein have not yet been carried out.

Here, we used deep mutational scanning to quantify the effects of thousands of NTD single mutations on S protein expression. One notable observation was that NTD residues, unlike RBD residues, showed a weak correlation between mutational tolerability and relative solvent accessibility (RSA). Instead, the mutational tolerability of NTD residues strongly correlated with their distance to RBD and S2. Residues S50 and G232 were two exceptions, in which they were proximal to S2 and RBD, respectively, and yet had a high mutational tolerability. Subsequently, we functionally characterized two mutations that increased S protein expression, namely S50Q and G232E. These results have important implications towards understanding NTD evolution and S-based immunogen design.

## RESULTS

### Most NTD mutations have minimal impact on S protein expression

To study how SARS-CoV-2 S protein expression is influenced by NTD mutations, we created a mutant library that contained all possible single amino acid mutations across residues 14-301 of the S protein. Each of these 288 residues was mutated with the choice of all 19 other amino acids and the stop codon, leading to a mutant library with 5,760 single amino acid mutations. The mutant library was expressed using the HEK293T landing pad cell system, such that each transfected cell stably expressed only one mutant^37, 38^. Fluorescence-activated cell sorting (FACS) was then performed using the human anti-S2 antibody CC40.8^39^, with PE anti-human IgG Fc as the secondary antibody. Four separated gates were set up based on the PE signals, each covering 25% of the entire population (**Figure S1**). The frequency of each mutant among the entire population was calculated (see Materials and Methods), and a cutoff of 0.0075% was set up to filter out mutants with potentially noisy measurements. Among the 5,760 missense and nonsense mutations, 3,999 (69%) of them satisfied the frequency cutoff for downstream analysis. Of note, the design of our mutant library adopted an internal barcoding strategy that uses synonymous mutations to facilitate sequencing error correction^40^. As described previously^41^, the expression score of each mutation was calculated based on their frequency in each of the four gates and normalized such that the average expression score of silent mutations was 1 and that of nonsense mutations was 0.

To evaluate the quality of the deep mutational scanning results, we assessed the expression score distributions of missense, nonsense, and silent mutations (**Figure S2A**). The difference between the expression scores of silent mutations and nonsense mutations was apparent and significant (P = 6×10^-166^), which validated the selectivity of the deep mutational scanning experiment. Interestingly, silent mutation and missense mutations had similar expression scores, although the difference is statistically significant (P = 2×10^-5^), indicating that most amino acid mutations in the NTD did not affect S protein expression. In addition, a Pearson correlation of 0.53 was obtained between the expression scores from two independent biological replicates (**Figure S2B**), demonstrating the reproducibility of the deep mutational scanning experiment.

To summarize the expression scores for individual mutations, a heatmap was generated (**Figure 1**). We noticed that high-expressing mutations were enriched within the five NTD loop regions (**Figure S3A**)^12^. High-expressing mutations were also found in residues outside of the loop regions, such as residues S50 and G232. This observation shows that some NTD mutations can improve the expression of S protein.

**Figure 1.**
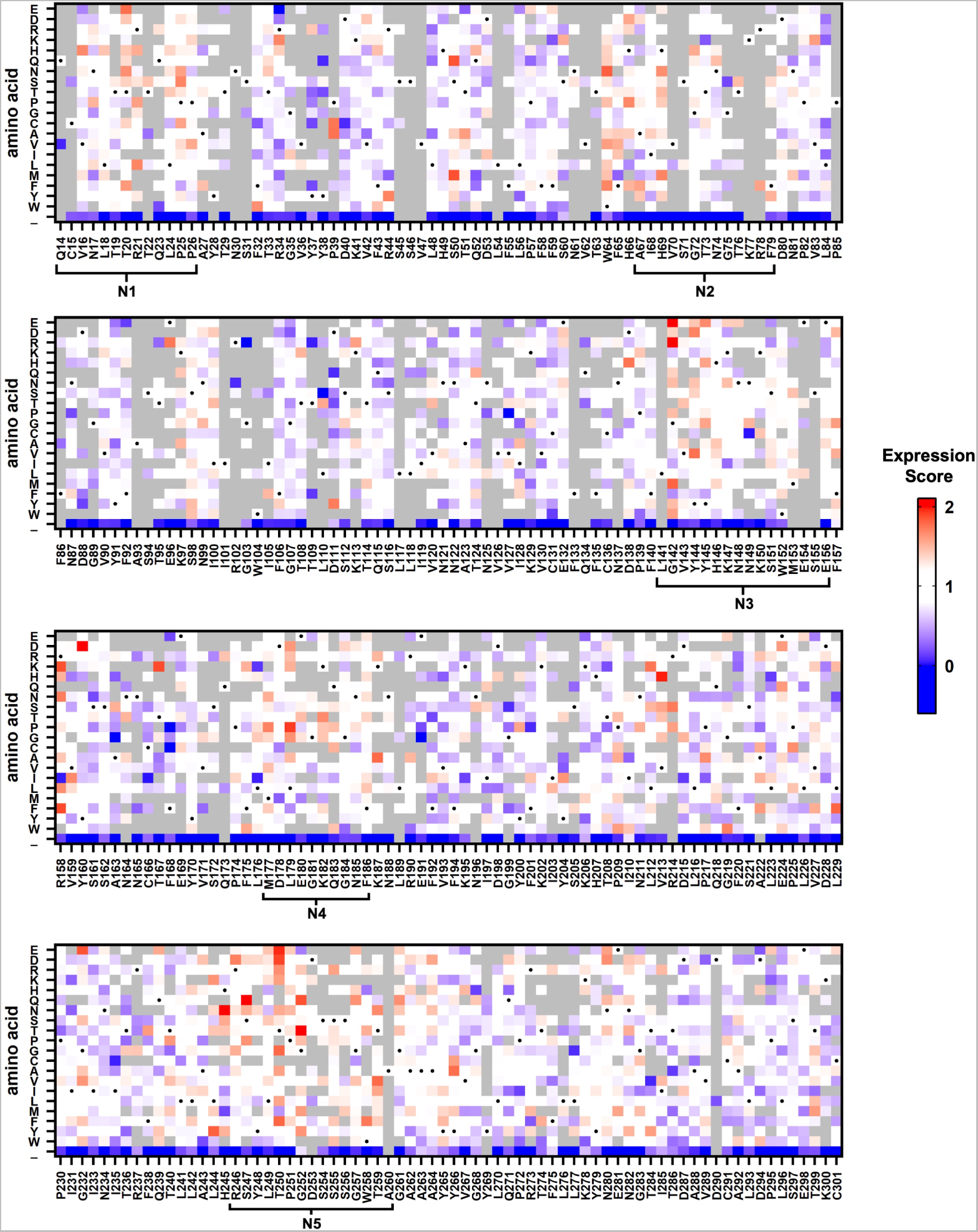
Effects of NTD single mutations on S protein expression. The expression scores of individual NTD mutations are shown as a heatmap. X-axis represents the residue position. Y-axis represents different amino acids as well as the stop codon (_). Amino acids corresponding to the WT sequence are indicated by the black dots. Mutations with a total frequency of <0.0075% were excluded from the analysis and shown in grey. Regions corresponding to the N1-N5 loops were defined as previously described^12^.

### Mutational tolerability has minimal correlation with solvent accessibility

While some residues were enriched in high-expression mutations (see above), others were enriched in low-expression mutations (e.g. residues D40, L84, and N234) (**Figure 1**). Consequently, we aimed to identify the biophysical determinants of mutational tolerability in terms of S protein expression. For each residue, we defined the mutational tolerability as the mean expression score of mutations. A higher mutational tolerability would indicate the enrichment of high-expressing mutations at the specified residue. In contrast, a lower mutational tolerability would indicate the enrichment of low-expressing mutations at the specified residue. A total of 243 NTD residues had six or more mutations with expression score available and were included in this analysis.

First, we investigated whether a correlation existed between the mutational tolerability and relative solvent accessibility (RSA). Since buried residues are typically important for protein folding stability, residues with a lower RSA are generally expected to have a lower mutational tolerability. For example, previous deep mutational scanning studies on the RBD have shown a decent correlation between RSA and mutational tolerability (Spearman correlation = 0.73, **Figure 2A**)^34, 42^. In contrast, the mutational tolerability of NTD residues had a much weaker correlation with RSA (Spearman correlation = 0.19, **Figure 2B**). These observations indicate that the folding stability of NTD does not have a strong influence on its mutational tolerability, and hence the S expression level.

**Figure 2.**
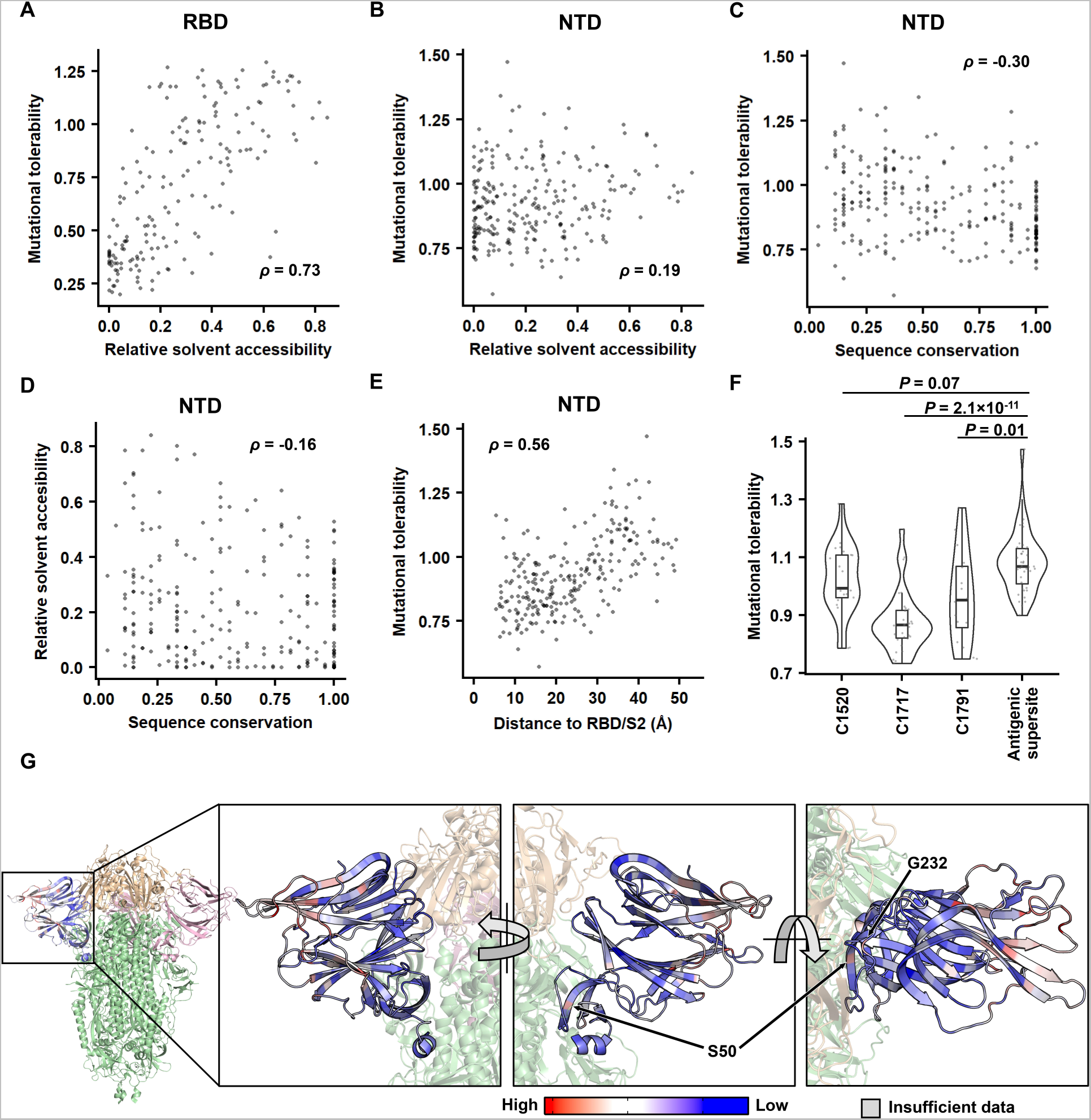
identifying the biophysical determinants of mutational tolerability. **(A-B)** The relationship between relative solvent accessibility (RSA) and the mutational tolerability is shown for **(A)** RBD and **(B)** NTD. The deep mutational scanning data on RBD expression was from a previous study^42^. **(C-D)** The relationship between sequence conservation among 27 sarbecovirus strains (**Table S6**) and **(C)** the mutational tolerability, or **(D)** RSA of each NTD residue is shown. **(E)** The relationship between the distance to RBD/S2 and the mutational tolerability of each NTD residue is shown. **(A-E)** Each datapoint represents one residue. The Spearman’s rank correlation coefficient (ρ) is indicated. **(F)** The mutational tolerability of residues within the cross-neutralizing NTD antibody epitopes (C1520, C1717, C1791)^16^ is compared to that within the antigenic supersite^14^ using a violin plot. Each datapoint represents one residue. P-values were computed by two-tailed t-test. **(G)** The mutational tolerability of each NTD residue is projected on one NTD of the S trimer structure (PDB 6ZGE^44^ and PDB 7B62^45^). Red indicates residues with higher mutational tolerability, while blue indicates residues with lower mutational tolerability. Residues with insufficient data to calculate mutational tolerability are colored in grey. Two residues of interests, namely S50 and G232, are indicated. RBDs are colored in wheat, the two other NTDs are in pink, and the rest of the S1 and S2 subunits are in green.

To investigate whether the mutational tolerability correlated with sequence conservation, we then analyzed the NTD sequences of 27 sarbecovirus strains, including SARS-CoV-2. Less conserved residues tended to have a higher mutational tolerability, while more conserved residues tended to have a lower mutational tolerability, although the correlation was not strong (Spearman correlation = -0.30) (**Figure 2C**). In comparison, the correlation between sequence conservation and RSA was even weaker (Spearman correlation = -0.16, **Figure 2D**).

### Mutational tolerability correlates with distance to RBD/S2

We further calculated the distance from each NTD residue to RBD/S2 of the S protein. A positive correlation was observed between the mutational tolerability and the distance to RBD/S2 (Spearman correlation = 0.56) (**Figure 2E**). In other words, the more distant an NTD residue was from the RBD/S2, the higher the mutational tolerability was. Such correlation was apparent when the mutational tolerability of each NTD residue was projected on the S protein structure **(Figure 2G)**. Consistently, the epitopes of two cross-neutralizing antibodies, namely C1717 and C1791, were significantly closer to RBD/S2 (P ≤ 5×10^-4^) and had lower mutational tolerability (P ≤ 0.01) when compared to the rapidly evolving NTD antigenic supersite (**Figure 2F and Figure S3B**)^14, 16, 43^.

### Two buried NTD mutations increase S protein expression

While NTD residues adjacent to RBD/S2 typically had a low mutational tolerability, S50 and G232 were two exceptions **(Figure 2G)**. For example, mutations S50G and G232E had a high expression score in our deep mutational scanning results. To validate this finding, we used the same landing pad system to construct HEK293T cell lines that stably expressed S50Q, G232E, and S50Q/G232E double mutant. As quantified by flow cytometry analysis (**Figure 3A and Figure S4**), the expression level of S50Q and G232E increased from wild type (WT) by 1.7-fold (P = 0.002) and 1.5-fold (P = 9 × 10^-4^), respectively, whereas that of S50Q/G232E increased by 2.5-fold (P = 2 × 10^-6^).

**Figure 3.**
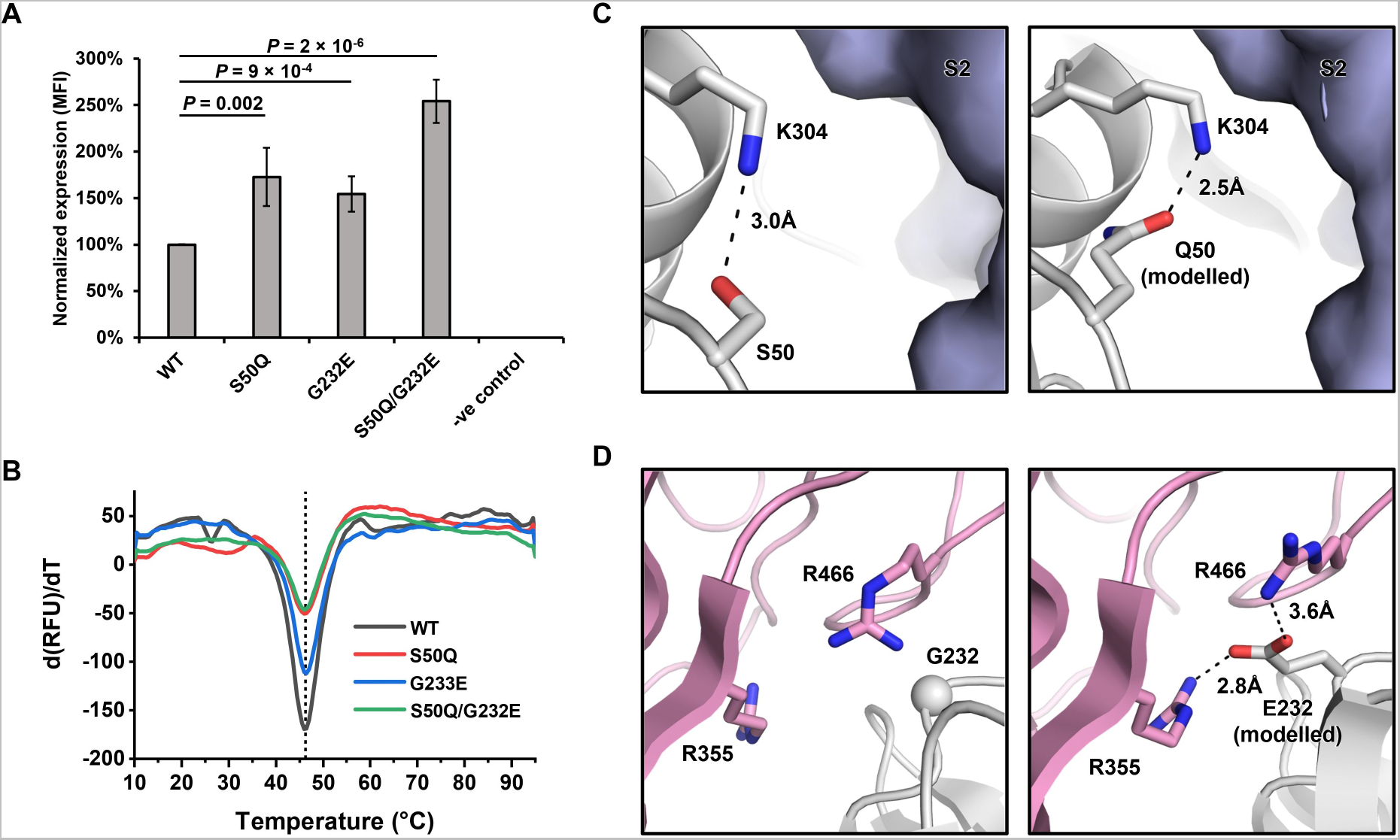
S50Q and G232E at the interdomain interface increase S protein expression. **(A)** Cell surface S protein expression of WT and NTD mutants was quantified using flow cytometry analysis with CC40.8 as the primary antibody. Untransfected HEK293T landing pad cells were used as a negative control (-ve control). S protein expression level was defined as the mean fluorescence intensity (MFI) of the positive gated population. S protein expression level was normalized to WT. The error bar indicates the standard deviation of six independent experiments. P-values were computed by two-tailed t-test. **(B)** Thermostability of WT S protein and selected NTD mutants was measured using differential scanning fluorimetry. The black vertical dotted line indicates the melting temperature of WT (T_m_ = 46.2 °C). **(C-D)** Rosetta-based structural modelling of **(C)** S50Q and **(D)** G232E was performed using the structure of S protein (PDB 6ZGE)^45^. The three protomers of the S protein are colored in white, light blue, and pink. Potential interactions are represented by black dashed lines with distance labeled.

To probe the structural impact of S50Q and G232E, we analyzed their local environments on the structure of S protein and performed structural modelling using Rosetta (**Figure 3C-D**)^44–46^. S50 forms a hydrogen bond with K304 and is proximal to the S2 subunit. Structural modelling showed that S50Q not only is able to maintain the hydrogen bond with K304, but also strengthens the van der Waals interaction between the NTD and S2 by pushing K304 towards S2 from the adjacent protomer (**Figure 3C**). G232 is proximal to a positively charged region on the RBD that is featured by R355 and R466 (**Figure 3D**). Structural modeling suggested that G232E could form favorable electrostatic interactions with both R355 and R466. We further recombinantly expressed these mutants and tested their thermostability using a thermal shift assay (**Figure 3B**). Of note, all the recombinantly expressed S proteins contained K986P/V987P mutations in the S2 subunit, which are known to stabilize the prefusion conformation and increase expression^26, 47^. The melting temperatures of WT and NTD mutants were almost identical at a T_m_ of 46 °C to 46.5 °C. These observations indicate that despite both S50Q and G232E improve the interaction between NTD and the rest of the S protein, they have minimal impact on the global folding stability of the S protein.

### S50Q and G232E have minimal effects on the fusion activity and antigenicity

To understand the functional consequences of S50Q and G232E, we further tested whether S50Q, G232E, and S50Q/G232E exhibited a change in fusion activity compared the WT. A fluorescence-based cell-cell fusion assay that relied on the split mNeonGreen2 (mNG2)^48^ was performed (see Materials and Methods, **Figure S5**). Briefly, HEK293T landing pad cells that expressed human ACE2 (hACE2) and mNG2_1-10_ were mixed with HEK293T landing pad cells that expressed S proteins and mNG2_11_. Green fluorescence due to mNG2 complementation was generated when fusion between the two cell lines occurred. Fluorescence microscopy analysis showed that all mutants facilitated hACE2-mediated fusion (**Figure 4A-B**). Consistently, flow cytometry analysis at both 3-hour and 24-hour post-mixing indicated that none of the tested mutants diminished the fusion activity when compared to WT (**Figure 4C-D**). At 3-hour post-mixing, both S50Q (24%, P = 0.03) and G232E (25%, P = 0.01) showed mild, yet significant, increases in fusion activity compared to WT. Similarly, at 24-hour post-mixing, S50Q (19%, P = 0.01), G232E (13%, P = 0.02), and S50Q/G232E double mutant (37%, P = 0.005) all showed an increase in fusion activity compared to WT. Such a mild increase in fusion activity may simply be attributed to the higher expression level of the mutants. Negative control cells expressing the K986P/V987P double mutant, which is known to stabilize the prefusion form of the S protein^26, 47^, did not show any fusion activity (**Figure 4A-D**).

**Figure 4.**
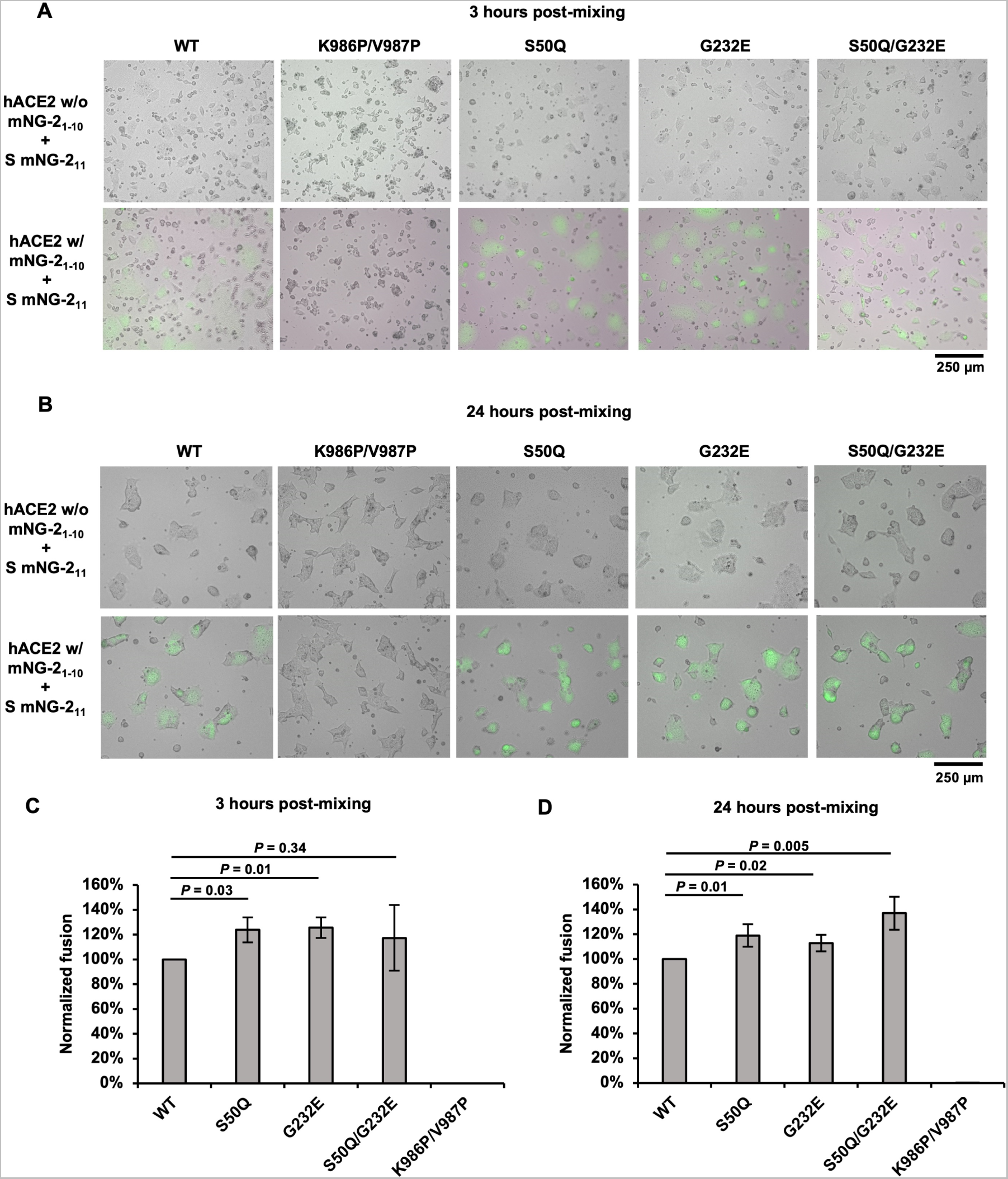
S50Q, G232E, and S50Q/G232E do not diminish fusion activity. **(A-B)** Fluorescence microscopy analysis of the fusion events at **(A)** 3-hour and **(B)** 24-hour post-mixing of cells expressing S and mNG2_11_ (S cells) and cells expressing hACE2 and mNG2_1-10_ (hACE2 cells). Cells with green fluorescence signals are the fused cells. Scale bars are shown at the bottom right corner. **(C-D)** Fusion activity of WT and selected NTD mutants at **(C)** 3-hour and **(D)** 24-hour post-mixing was quantified using flow cytometry analysis. Fusion activity was normalized to WT. The error bar indicates the standard deviation of at least four independent experiments. P-values were computed by two-tailed t-test.

We then proceeded to investigate whether S50Q, G232E, and S50Q/G232E alter the antigenicity of the S protein. The binding of three antibodies targeting different domains of the S protein were tested, namely CC12.3 (anti-RBD)^49^, S2M28 (anti-NTD)^14^, and COVA1-07 (anti-S2)^50^. Flow cytometry analysis showed that all three antibodies bound to the tested mutants at a similar level as WT (**Figure 5 and Figure S6**), indicating that S50Q, G232E, and S50Q/G232E did not alter the structural conformation and antigenicity of the S protein.

**Figure 5.**
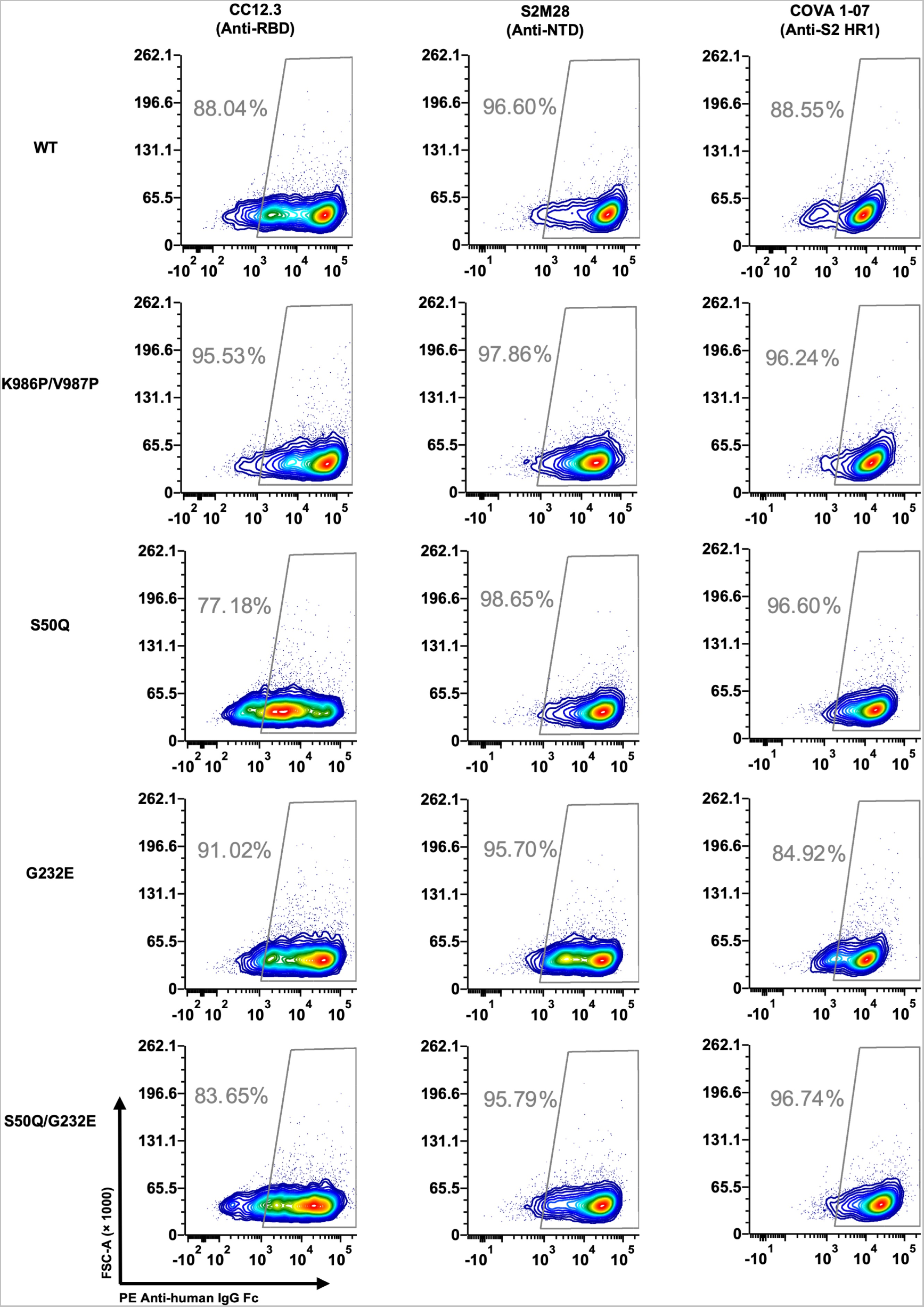
S50Q, G232E, and S50Q/G232E do not alter antibody binding. Three antibodies targeting different domains on the S were tested for binding to cells expressing WT, K986P/V987P, S50Q, G232E, or S50Q/G232E S protein. Binding was measured by flow cytometry analysis. Gating was set up using untransfected HEK293T landing pad cells, which served as a negative control (**Figure S6**).

## DISCUSSION

S protein is central to the research of SARS-CoV-2 evolution and COVID-19 vaccines^51–54^. While both the RBD and the NTD on the S protein are targets of neutralizing antibodies and involve in the antigenic drift of SARS-CoV-2^43, 55–61^, the NTD often receives less attention than the RBD. Using deep mutational scanning, this study shows that many NTD mutations at buried residues do not affect S protein expression. At the same time, the closer an NTD mutation is to RBD/S2, the more likely it is detrimental to S protein expression. These observations imply that for optimum S protein expression, the structural stability at the NTD-RBD and the NTD-S2 interfaces is more critical than the folding stability of the NTD. Our results also at least partly explain why the N1 to N5 loops, which contain the NTD antigenic supersite^62^ and are far from the NTD-RBD/S2 interfaces, are highly diverse among SARS-CoV-2 variants and sarbecovirus strains. Overall, this study provides crucial biophysical insights into the evolution of the NTD.

NTD mutations S50Q and G232E, which locate at the interdomain interface and increase S protein expression, represent another important finding of this study. Engineering high expressing S protein can lower the production cost of recombinant COVID-19 vaccine and may improve the effectiveness of mRNA vaccines^25^. Similar to certain previously characterized mutations in the S2^26, 27^, S50Q and G232E in the NTD increase the expression yield of the S protein without changing its T_m_. Consistently, a recent study showed that NTD mutations in BA.1 improve the expression of S protein without increasing its thermostability^63^. Furthermore, S50Q and G232E are not solvent exposed on the S protein surface and do not seem to alter the antigenicity of the S protein. Of note, according to our deep mutational scanning data, S50Q and G232E are just two of many mutations that enhance S protein expression. Therefore, although most studies on S-based immunogen design focus on the mutations in the RBD and S2^7, 26–29^, our results suggest that mutations in NTD can provide a complementary strategy.

We acknowledge that S protein expression level does not necessarily correlate with virus replication fitness. For example, NTD mutations that do not affect the S protein expression may be detrimental to the replication fitness of SARS-CoV-2, due to negative impact on NTD functionality. While the functional importance of the NTD in natural infection remains largely unclear, NTD has been proposed to facilitate virus entry by interacting with DC-SIGN, L/SIGN, AXL, ASGR1, and KREMEN1^64–66^. Studies have also shown that the NTD can allosterically evade antibody binding by interacting with a heme metabolite^45^, as well as modulate the efficiency of virus-host membrane fusion^67, 68^. To fully comprehend the biophysical constraints of NTD, future studies should systematically investigate how different NTD mutations affect virus replication fitness.

## MATERIALS AND METHODS

### Construction of the NTD mutant library

SARS-CoV-2 S NTD mutant library was constructed based on the HEK293T landing pad system^37, 38^. The template for constructing the NTD mutant library was a plasmid that encoded (from 5’ to 3’) an attB site, a codon-optimized SARS-CoV-2 S (GenBank ID: NC_045512.2) with the PRRA motif in the furin cleavage site deleted, an internal ribosome entry site (IRES), and a puromycin-resistance marker. This plasmid was used as a PCR template to generate a linearized vector and a library of mutant NTD inserts. The linearized vector was generated using 5’-TGC TCG TCT CTA CAA CTC CGC CAG CTT CAG CAC C-3’ and 5’-TGC TCG TCT CTT CAC TGG CCG TCG TTT TAC AAC G-3’ as primers. Inserts were generated by two separate batches of PCRs to cover the entire NTD. The first batch of PCRs consisted of 36 reactions, each containing one cassette of forward primers as well as the universal reverse primer 5’-TGC TCG TCT CGT TGT ACA GCA CGG AGT AGT CGG C-3’. Each cassette contained an equal molar ratio of eight forward primers that had the same 21 nt at the 5’ end and 15 nt at the 3’ end. Each primer within a cassette were also encoded with an NNK (N: A, C, G, T; K: G, T) sequence at a specified codon positions for saturation mutagenesis. In addition, each primer also carried unique silent mutations (also known as synonymous mutations) to help distinguish between sequencing errors and true mutations in downstream sequencing data analysis as described previously^40^. The forward primers, named as CassetteX_N (X: cassette number, N: primer number), are listed in **Table S1**. The second batch of PCR consisted of another 36 PCRs, each with a universal forward primer 5’-TGC TCG TCT CAG TGA ATT GTA ATA CGA CTC ACT A-3’ and a unique reverse primer as listed in **Table S2**. Subsequently, 36 overlapping PCRs were performed using the universal forward and reverse primers, as well as a mixture of 10 ng each of the corresponding products from the first and second batches of PCR. The 36 overlap PCR products were then mixed at equal molar ratio to generate the final insert of the NTD mutant library. All PCRs were performed using PrimeSTAR Max polymerase (Takara Bio) per manufacturer’s instruction, followed by purification using Monarch Gel Extraction Kit (New England Biolabs). The final insert and the linearized vector were digested by BsmBI-v2 (New England Biolabs) and ligated using T4 DNA Ligase (New England Biolabs). Ligation product was purified by PureLink PCR Purification Kit (Thermo Fisher Scientific) and then transformed into MegaX Dh10B T1R cells (Thermo Fisher Scientific). At least half a million colonies were collected. Plasmid mutant library were purified from the bacteria colonies using PureLink HiPure Plasmid Midiprep Kit (Invitrogen). All primers in this study were ordered from Integrated DNA Technologies.

### Construction of stable cell lines using HEK293T landing pad cells

Human embryonic kidney 293T (HEK293T) landing pad cells^37, 38^ were used to display the NTD mutant library for deep mutational scanning. Landing pad cells were maintained using complete growth medium consisting of Dulbecco’s Modified Eagle’s Medium (DMEM) (Corning), 10% v/v FBS (VWR), Pen-Strep (Gibco), non-essential amino acid (Gibco), and 2 μg/mL doxycycline. 1.2 μg of plasmid was transfected into 6 × 10^5^ landing pad cells. For the deep mutational scanning experiment, eight transfection reactions were carried out in parallel to minimize loss of mutant diversity at the transfection step. Transfected cells were then incubated at 37 °C with 5% CO_2_. After 48 hours, 10 nM AP1903 was supplemented to carry out negative selection. At 72 hours after the negative selection, positive selection antibiotic (1 μg/mL puromycin for NTD cell lines or 100 μg/mL hygromycin for hACE2 cell lines) was supplemented to the medium to carry out positive enrichment of cells with successful recombination. Constructed cell lines would remain in the complete growth medium supplemented with doxycycline and the positive selection antibiotics.

### Sorting the NTD mutant library based on S protein expression level

Four T-75 flasks (Corning) that were 90% confluent with cells that carried the NTD mutant library were washed with 1× PBS, harvested with warm versene and pelleted via centrifugation at 300 × *g* for 5 mins at room temperature. Cells were then resuspended in FACS buffer (2% v/v fetal bovine serum, 5 mM EDTA in DMEM supplemented with glucose, L-glutamine and HEPES but without phenol red (Gibco)). Subsequently, cells were incubated with 5 μg/mL of CC40.8 at 4 °C with gentle shaking for 1 hour. Cells were washed once with ice-cold FACS buffer and incubated with 1 μg/mL PE anti-human IgG Fc (Biolegend) at 4 °C with gentle shaking in the dark for 1 hour. Cells were washed once and resuspended in ice-cold FACS buffer. Cells were then filtered using a 40 μm cell strainer (VWR) before cell sorting. FACS were performed using a BD FACSAria II cell sorter (BD) with a 561 nm laser and a 582/15 bandpass filter. Cells were collected into ice-cold D10 medium (Dulbecco’s modified Eagle medium with 4.5 g/L glucose, 4 mM L-glutamine and 110 mg/L sodium pyruvate (Corning), supplemented with 10% v/v fetal bovine serum (VWR), 1× penicillin-streptomycin (Gibco), 1× non-essential amino acids (Gibco)) and binned into no (bin 0), low (bin 1), medium (bin 2) and high (bin 3) expression according to PE signal, where each bin contains 25% of the singlet population (**Figure S1**). A biological replicate of the deep mutational scanning experiment was performed, starting from the transfection step.

### Next-generation sequencing of the NTD mutant library

Sorted cells from each bin were pelleted at 300 × g, 4 °C for 15 mins and then resuspended in 200 μL PBS (Corning). Genomic DNA extraction was then performed using DNA Blood and Tissue Kit (QIAGEN) according to the manufacturer’s instructions with a modification: cells were incubated at 56 °C for 30 min instead of 10 min. The NTD mutant library was amplified from the genomic DNA in two non-overlapping fragments using KOD DNA polymerase (MilliporeSigma) per manufacturer’s instruction with the following two primer sets, respectively (also see **Table S3**). Set 1: 5’-CAC TCT TTC CCT ACA CGA CGC TCT TCC GAT CTC TGC TGC CTC TGG TGT CCA GC-3’ (NTD-DMS-recover-1F) and 5’-GAC TGG AGT TCA GAC GTG TGC TCT TCC GAT CTG TTG GCG CTG CTG TAC ACC CG-3’ (NTD-DMS-recover-1R) Set 2: 5’-CAC TCT TTC CCT ACA CGA CGC TCT TCC GAT CTA GCT GGA TGG AAA GCG AGT TC-3’ (NTD-DMS-recover-2F) and 5’-GAC TGG AGT TCA GAC GTG TGC TCT TCC GAT CTC ACG GTG AAG GAC TTC AGG GT-3’ (NTD-DMS-recover-2R) A second round of PCR was carried out to add the adapter sequence and index to the amplicons as described previously^69^. The final PCR products were submitted for next-generation sequencing using Illumina MiSeq PE300.

### Analysis of next-generation sequencing data

Next-generation sequencing data were obtained in FASTQ format. Forward and reverse reads of each paired-end read were merged by PEAR^70^. The merged reads were parsed by SeqIO module in BioPython^71^. Primer sequences were trimmed from the merged reads. Trimmed reads with lengths inconsistent with the expected length were discarded. The trimmed reads were then translated to amino acid sequences, with sequencing error correction performed at the same time as previously described^40^. Amino acid mutations were called by comparing the translated reads to the WT amino acid sequence. Frequency (*F*) of a mutant *i* at position *s* within bin *n* of replicate *k* was computed for each replicate as follows:

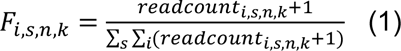

A pseudocount of 1 was added to the read counts of each mutant to avoid division by zero in subsequent steps. We then calculated the total frequency (*F_total_*) of mutant *i* at position *s* as follows:

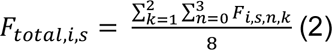

Mutants with a *F_total_* of equal or greater than 0.0075% were selected for downstream analysis. Subsequently, the weighted average (*W*) of each mutant among 4 bins (bin 0 to bin 3) in each replicate was computed as described previously^41^:

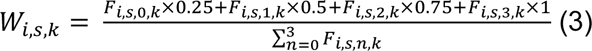

Selected mutants were then categorized based on the mutation types (missense, nonsense, and silent). The mean value of weighted average for nonsense as well as silent mutations were calculated. Expression score (*ES*) of a mutant *i* at position s of replicate *k* was calculated as described previously^41^:

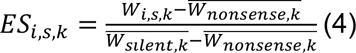

Final expression score of a mutant *i* at position *s* was calculated by taking the average of the expression scores between replicates. Mutational tolerability of position *s* was then calculated by taking the average of the expression scores of all mutants at that position:

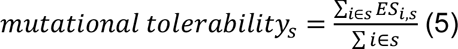

### Structural analysis of deep mutational scanning results

DSSP^72, 73^ was used to calculate the solvent exposure surface area (SASA) of each residue in NTD and RBD on the S trimer (PDB 6ZGE)^44^. Deep mutational scanning result of RBD was extracted from a previous study^42^. Relative solvent accessibility (RSA) was computed by dividing the SASA by the theoretical maximum allowed solvent accessibility of the corresponding amino acid^74^.

Each NTD residue’s distance to RBD/S2 was calculated based on the S trimer structure (PDB 6ZGE)^44^ with the NTD replaced by the high resolution crystal structure (PDB 7B62)^45^. For each NTD residue, the distances to all RBD and S2 residues were measured. The shortest distance was then recorded as the “distance to RBD/S2”. Residue-residue distance was defined as the distance between the centroid coordinates of two residues.

To visualize the mutational tolerability of each NTD residue, the crystal structure of SARS-CoV-2 S protein NTD (PDB 7B62) was used^45^. The NTD crystal structure was then aligned with the S trimer to generate the figures (PDB 6ZGE)^44^.

### NTD sequence conservation analysis

The sequence conservation analysis of NTD was based on 27 sarbecovirus strains (**Table S6**)^1, 75–79^. S sequences of these stains were retrieved from GenBank and Global Initiative for Sharing Avian Influenza Data (GISAID)^80^. Their NTD sequences were then identified using tBlastn search using the amino acid sequence of SARS-CoV-2 Hu-1 NTD (Gene ID: 43740568) as the query sequence. The BlastXML output of the tBlastn was then parsed and used as the input for multiple sequence alignment using MAFFT^81, 82^. For each residue position, sequence conservation was defined as the proportion of strains that contains the same amino acid variant as SARS-CoV-2 Hu-1.

### Rosetta-based mutagenesis

The structure of the spike protein was obtained from Protein Data Bank (PDB 6ZGE)^44^. Water molecules and N-acetyl-D-glucosamine were removed using PyMOL (Schrödinger). Then, the amino acids were renumbered using pdb-tools^83^. Fixed backbone point-mutagenesis for S50Q and G232E was performed using the ‘fixbb’ application in Rosetta (RosettaCommons). One-hundred poses were generated for each mutagenesis. Using the lowest-scoring structure from fixed backbone mutagenesis as input, a constraint file was obtained using the minimize_with_cst application in Rosetta. Fast relax was then performed via the ‘relax’ application in Rosetta^46^ with the corresponding constraint file. The lowest-scoring structure out of eight was then used for structural analysis. Code and source files for structural modelling are available in https://github.com/nicwulab/SARS-CoV-2_NTD_DMS/tree/main/rosetta.

### Split mNeonGreen2-based cell-cell fusion assay

Human ACE2 (hACE2) construct was constructed in a previous study^38^. A split mNeonGreen2 (mNG2) reporter system was integrated into the S plasmid (see above) and the hACE2 plasmid^48^. Specifically, a gene fragment that encoded (from 5’ to 3’) a GCN4 leucine zipper, a GS linker, mNG2_1-10_, and a 2A self-cleaving peptide was inserted into the hACE2 plasmid between the IRES and the hygromycin resistance marker. Similarly, a gene fragment that encodes (from 5’ to 3’) a GCN4 leucine zipper, a GS linker, mNG2_11_, and a 2A self-cleaving peptide was inserted into the S plasmid between the IRES and the puromycin resistance marker. Each plasmid construct was transfected and recombined into HEK293T landing pad cells per steps described above.

Once the stable cell lines were created, 5 × 10^5^ landing pad cells expressing hACE2 with mNG2_1-10_ were seeded in 6-well plates (Fisher Scientific). The cells were then then incubated at 37 °C with 5% CO_2_ for 15 mins to allow seeding. Subsequently, 5 × 10^5^ landing pad cells expressing the S with mNG2_11_ were then added dropwise to the seeded hACE2 cells. Both cells were filtered through 40 μm cell strainer (VWR) prior to seeding. At 3-hour and 24-hour post-mixing, fusion events in each well were qualitatively assessed with an ECHO Revolve epifluorescence microscope (ECHO) in inverted format. Overlayed images were captured on white light and FITC filter channels using an UPlanFL N 10X/0.30NA objective (Olympus) with identical light intensity and exposure settings for all conditions. Cells in each well were then collected using 0.5 mM EDTA, pelleted via centrifugation at 300 × *g* for 5 mins at room temperature, and resuspended in the FACS buffer. LSRII flow cytometry (BD) was used to quantify the fusion events of each sample. Negative controls were measured first to set up proper gating strategies (**Figure S5**). Then, the flow cytometry analysis was performed on 10^5^ live cells for each sample. Data were analyzed using FCS Express 6 software (De Novo Software). The percentage of mNG2 positive population of each sample were used for normalization (**Table S4**).

### Flow cytometry analysis for the protein expression assay and antibody binding assay

Approximately 1 × 10^6^ cells that carried the selected SARS-CoV-2 S NTD mutant were washed with 1× PBS, harvested with warm versene and pelleted via centrifugation at 300 × g for 5 mins at room temperature. The cells were resuspended in the FACS buffer. Subsequently, cells were incubated with 5 μg/mL of the selected antibodies at 4 °C with gentle shaking for 1 hour. Cells were then washed once with ice-cold FACS buffer and incubated with 2 μg/mL PE anti-human IgG Fc (BioLegend) at 4 °C with gentle shaking in the dark for 1 hour. Cells were washed once, pelleted via centrifugation at 300 × g for 5 mins at room temperature, and resuspended in ice-cold FACS buffer. LSRII flow cytometry (BD) was used to measure the PE signal of each sample. Negative controls were measured first to set up proper gating strategies (**Figure S4 and S6**). Then, the flow cytometry analysis was performed on 10^5^ singlets for each sample. Data were analyzed using FCS Express 6 software (De Novo Software).

### Normalization of the expression assay results

The mean fluorescence intensity (MFI) of the entire population was recorded for each sample, followed by the normalization as previously described^42^. For a given sample *i*, the following equation was used to compute the normalized expression (*NE*):

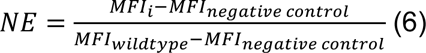

Normalizations were performed for each sample within a given biological replicate (**Table S4**).

### Recombinant expression and purification of soluble S protein

SARS-CoV-2 S ectodomain with the PRRA motif in the furin cleavage site deleted and mutations K986P/V987P, which are known to stabilize the prefusion conformation and increase expression^26, 47^, was cloned into a phCMV3 vector. The S ectodomain construct contained a trimerization domain and a 6×His-tag at the C-terminal. Expi293F cells (Gibco), which were maintained using Expi293 expression medium (Gibco), were used to express soluble S protein. Briefly, 20 μg of the plasmid was transfected into 20 mL of Expi293F cells at 3 × 10^6^ cells mL^-1^ using ExpiFectamine 293 Transfection Kit (Thermo Fisher Scientific) following the manufacturer’s instructions. Transfected cells were then incubated at 37 °C, 8% CO_2_ and shaking at 125 rpm for 6 days. Cell cultures were then harvested and centrifuged at 4000 × *g* at 4 °C for 15 mins. The supernatant was clarified using a 0.22 μm polyethersulfone filter (Millipore). S protein in the clarified supernatant was then purified using Nickel Sepharose Excel resin (Cytiva), with 20 mM imidazole in PBS as wash buffer, and 300 mM imidazole in PBS as elution buffer. Three rounds of 2 mL elutions were performed. The eluted protein was then concentrated and analyzed by SDS-PAGE reducing gels (Bio-Rad) (**Figure S7A**). Concentrated protein solution was further purified using Superdex 200 XK 16/100 size exclusion column (Cytiva) in 20 mM Tris-HCl pH 8.0 and 150 mM NaCl (**Figure S7B**). Selected elution fractions were combined and concentrated. Final protein concentration was measured using NanoDrop One (Thermo Fisher Scientific).

### Protein thermostability assay

5 μg of purified protein was mixed with 5× SYPRO orange (Thermo Fisher Scientific) in 20 mM Tris-HCl pH 8.0, 150 mM NaCl at a final volume of 25 µL. The sample mixture was then transferred into an optically clear PCR tube (VWR). SYPRO orange fluorescence data in relative fluorescence unit (RFU) was collected from 10 °C to 95 °C using CFX Connect Real-Time PCR Detection System (Bio-Rad). The temperature corresponding to the lowest point of the first derivative, −d(RFU)/dT, was defined as the melting temperature (T_m_). Data were analyzed using OriginPro 2020b (Origin Lab). Raw data are shown in **Table S5**.

### Code availability

Custom python and R scripts for data analysis and plotting in this study have been deposited to https://github.com/nicwulab/SARS-CoV-2_NTD_DMS.

### Data availability

Raw sequencing data have been submitted to the NIH Short Read Archive under accession number: BioProject PRJNA792013. Biological materials including plasmids, constructed NTD mutant library as well as individual HEK293T landing pad cell lines are available by contacting the corresponding author (N.C.W.). Source data are provided with this paper.

## ACKNOWLEDGEMENT

We thank Meng Yuan, Huibin Lv, and Qi Wen Teo for helpful discussion and the Roy J. Carver Biotechnology Center at the University of Illinois at Urbana-Champaign for assistance with fluorescence-activated cell sorting and next-generation sequencing. This work was supported by National Institutes of Health (NIH) R00 AI139445 (N.C.W.), DP2 AT011966 (N.C.W.), R01 AI167910 (N.C.W.), the Michelson Prizes for Human Immunology and Vaccine Research (N.C.W.).

## AUTHOR CONTRIBUTIONS

W.O.O, T.J.C.T., and N.C.W. conceived and designed the study. G.S. and R.A. provided the CC40.8 antibody. K.A.M. provided the HEK293T landing pad cell line and assisted in experimental design. W.O.O and T.J.C.T. performed the deep mutational scanning and flow cytometry experiments. T.J.C.T. performed structural modelling using Rosetta. W.O.O, T.J.C.T., and R.L. expressed and purified the recombinant proteins. W.O.O, T.J.C.T., and C.K. performed the microscopy analysis. W.O.O, T.J.C.T., and N.C.W. performed data analysis. W.O.O., T.J.C.T. and N.C.W. wrote the paper and all authors reviewed and/or edited the paper.

