## Supplementary Figures (S1-S7) for "Probing the biophysical constraints of SARS-CoV-2 spike N-terminal domain using deep mutational scanning"

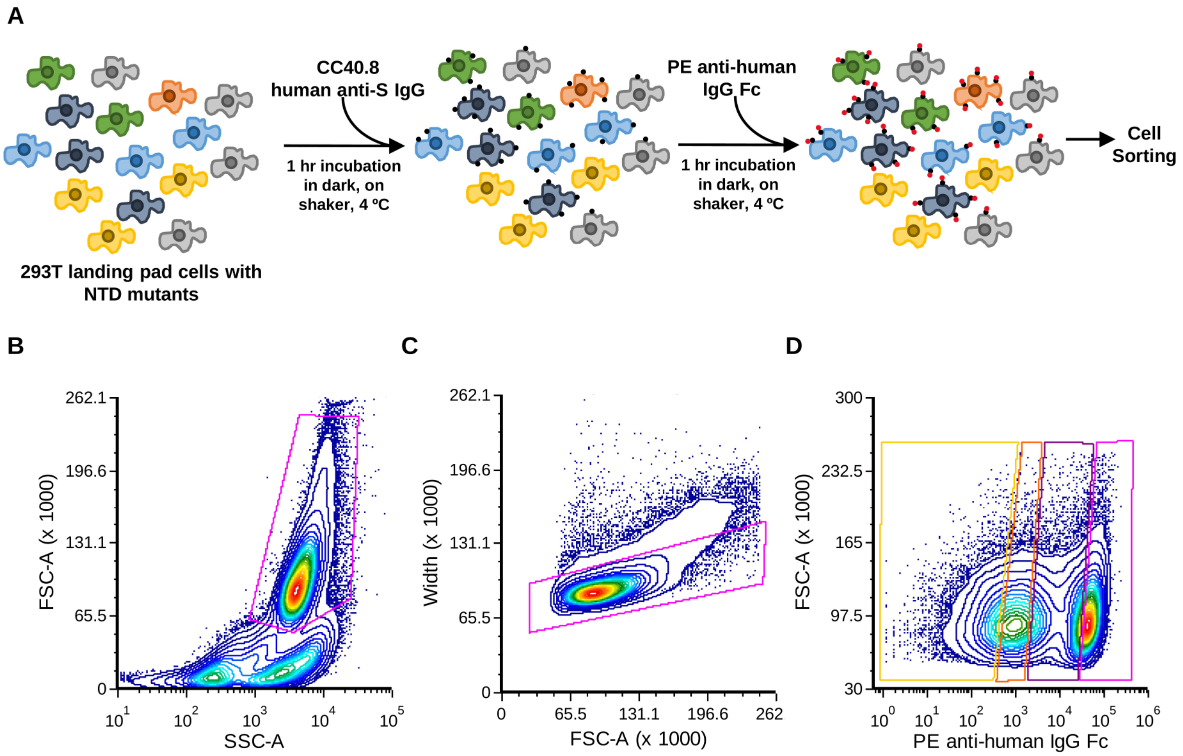


E

| Population | Replicate 1 (# of cells) | Replicate 2 (# of cells) |
|---|---|---|
| Bin 0 (PE-) | $2.303 \times 10^6$ | $2.356 \times 10^6$ |
| Bin 1 (PE+) | $2.351 \times 10^6$ | $2.324 \times 10^6$ |
| Bin 2 (PE++) | $2.372 \times 10^6$ | $2.333 \times 10^6$ |
| Bin 3 (PE+++) | $2.450 \times 10^6$ | $2.328 \times 10^6$ |

**Supplementary Figure 1. Overview of the NTD deep mutational scanning workflow. (A)** Schematics of NTD deep mutational scanning (see Materials and Methods for details). The black and red dots represent CC40.8 and PE anti-human IgG Fc, respectively. **(B-D)** Gating strategy for FACS is shown. **(B)** Live cells were first gated, then **(C)** the singlets among the live cells were gated, then **(D)** the singlets were sorted into four bins based on the PE signals, each covering 25% of the singlet population. **(E)** Sorting statistics of FACS. Each bin is color coded to correspond each of the four bins shown in **(D)**.

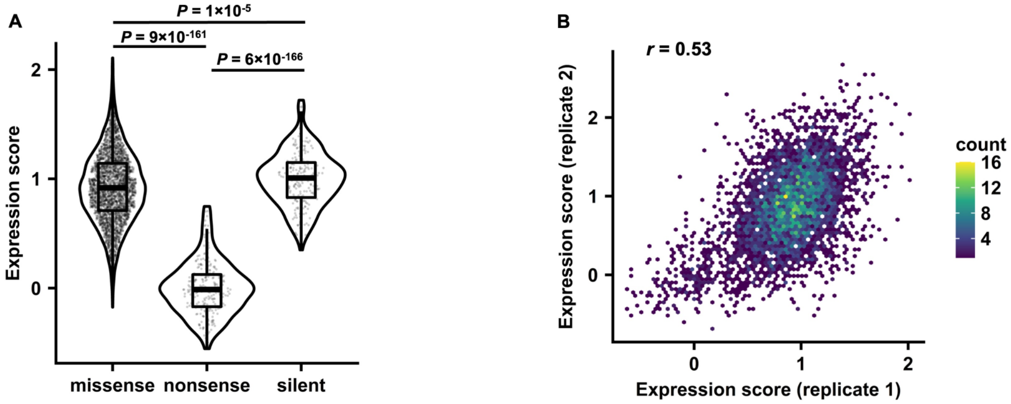


**Supplementary Figure 2. Data quality and reproducibility of the NTD deep mutational scanning experiment. (A)** The expression score distributions for missense, nonsense, silent mutations are shown as a violin plot. Each datapoint represents one mutation. P-values were computed by two-tailed t-test. **(B)** Correlation of expression scores between two biological replicates is shown as a density scatterplot. The Pearson correlation coefficient (r) is indicated.

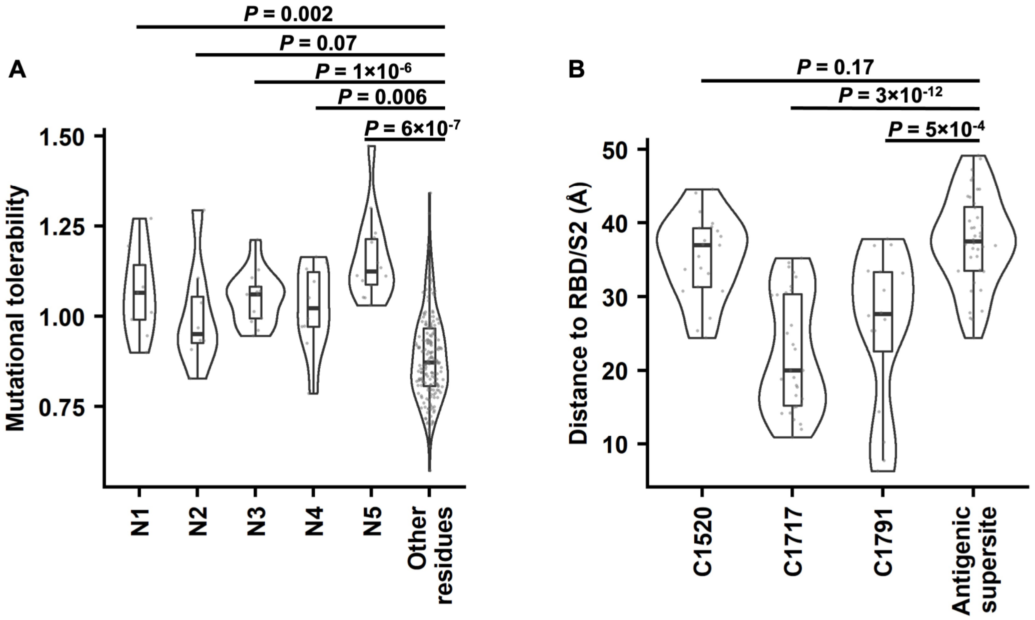


**Supplementary Figure 3. Characterization of selected regions on NTD. (A)** The difference in mutational tolerability between NTD loop regions (N1-N5) and other NTD residues is illustrated by a violin plot. Each datapoint represents one residue. Regions corresponding to the N1-N5 loops were defined as previously described[12]. **(B)** The difference in distance to RBD/S2 between epitopes of three cross-neutralizing antibodies[16] and the antigenic supersite[14] is illustrated by a violin plot. Each datapoint represents one residue. P-values were computed by two-tailed t-test.

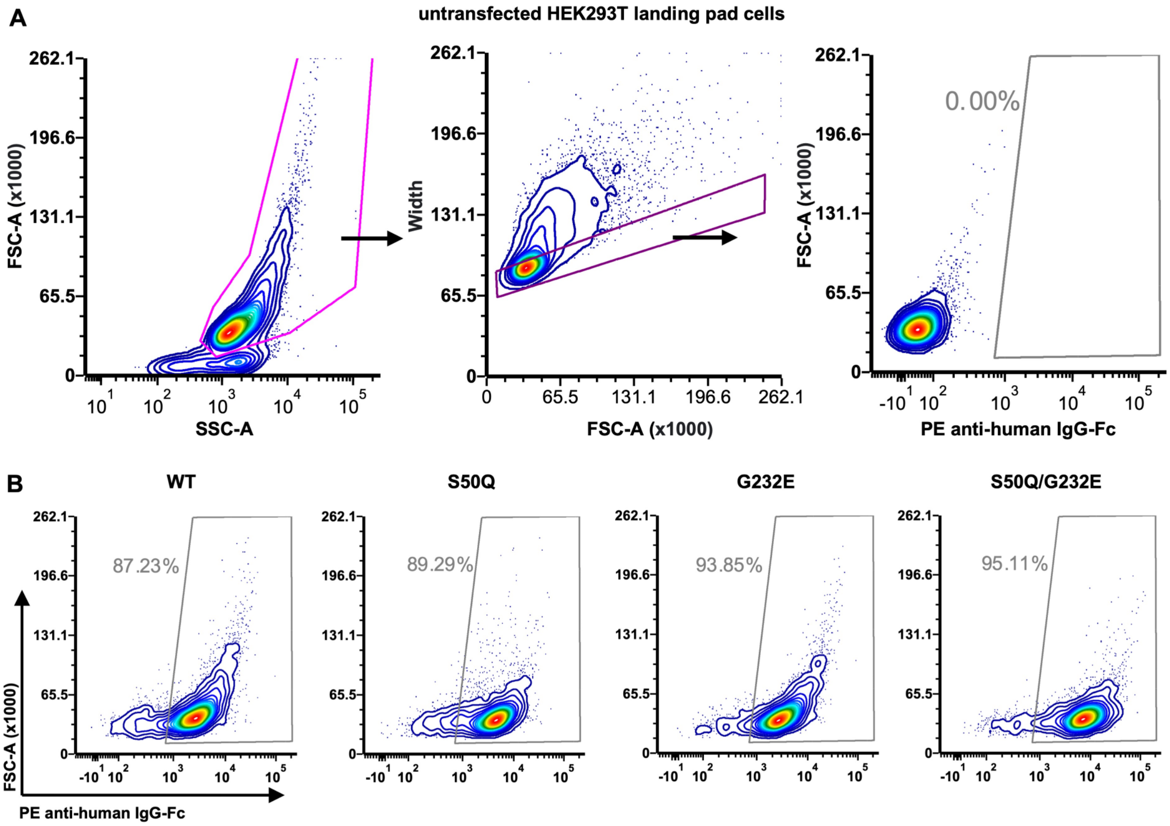


**Supplementary Figure 4. Flow cytometry analysis of S protein expression. (A)** Gating strategy for measuring the cell surface expression level of WT and mutant S proteins was setup based on untransfected HEK293T landing pad cells, which serve as a negative control. **(B)** Representative flow cytometry results of the WT and mutants.

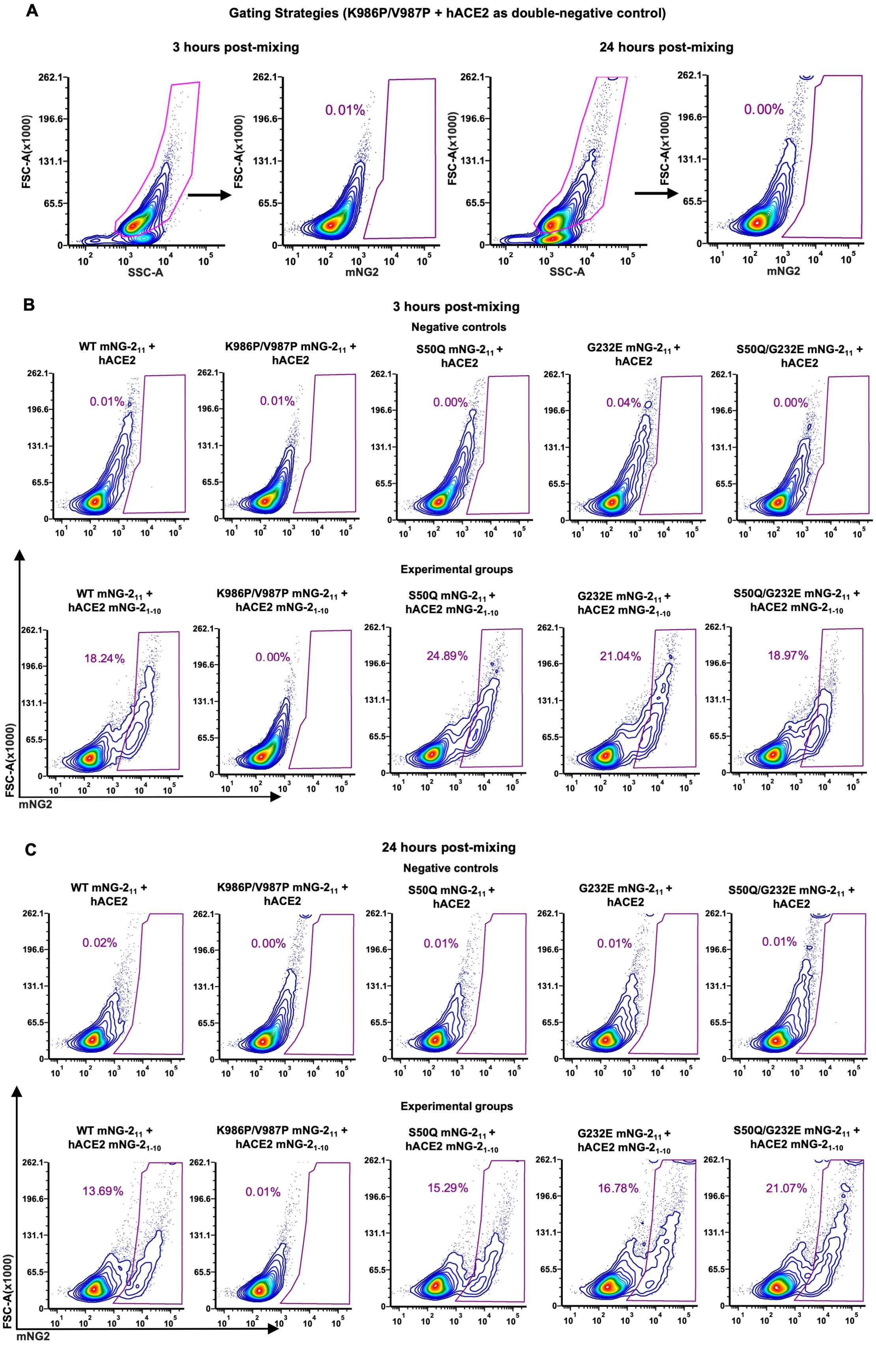
A
Gating Strategies (K986P/V987P + hACE2 as double-negative control)
3 hours post-mixing
24 hours post-mixing
FSC-A(x1000)
SSC-A
mNG2
0.01%
0.00%
B
3 hours post-mixing
Negative controls
WT mNG-2$_{11}$ + hACE2
K986P/V987P mNG-2$_{11}$ + hACE2
S50Q mNG-2$_{11}$ + hACE2
G232E mNG-2$_{11}$ + hACE2
S50Q/G232E mNG-2$_{11}$ + hACE2
0.01%
0.01%
0.00%
0.04%
0.00%
Experimental groups
WT mNG-2$_{11}$ + hACE2 mNG-2$_{1-10}$
K986P/V987P mNG-2$_{11}$ + hACE2 mNG-2$_{1-10}$
S50Q mNG-2$_{11}$ + hACE2 mNG-2$_{1-10}$
G232E mNG-2$_{11}$ + hACE2 mNG-2$_{1-10}$
S50Q/G232E mNG-2$_{11}$ + hACE2 mNG-2$_{1-10}$
18.24%
0.00%
24.89%
21.04%
18.97%
C
24 hours post-mixing
Negative controls
WT mNG-2$_{11}$ + hACE2
K986P/V987P mNG-2$_{11}$ + hACE2
S50Q mNG-2$_{11}$ + hACE2
G232E mNG-2$_{11}$ + hACE2
S50Q/G232E mNG-2$_{11}$ + hACE2
0.02%
0.00%
0.01%
0.01%
0.01%
Experimental groups
WT mNG-2$_{11}$ + hACE2 mNG-2$_{1-10}$
K986P/V987P mNG-2$_{11}$ + hACE2 mNG-2$_{1-10}$
S50Q mNG-2$_{11}$ + hACE2 mNG-2$_{1-10}$
G232E mNG-2$_{11}$ + hACE2 mNG-2$_{1-10}$
S50Q/G232E mNG-2$_{11}$ + hACE2 mNG-2$_{1-10}$
13.69%
0.01%
15.29%
16.78%
21.07%

**Supplementary Figure 5. Flow cytometry data evaluating the fusion activity of NTD mutants.** **(A)** Gating strategy for the fusion assay was setup based on the mixture of cells that express K986P/V987P S protein (without $mNG2_{11}$) and cells that express hACE2 (without $mNG2_{1\text{-}10}$). **(B-C)** Representative flow cytometry results for the fusion assay at **(B)** 3-hour post-mixing, and **(C)** 24-hour post-mixing.

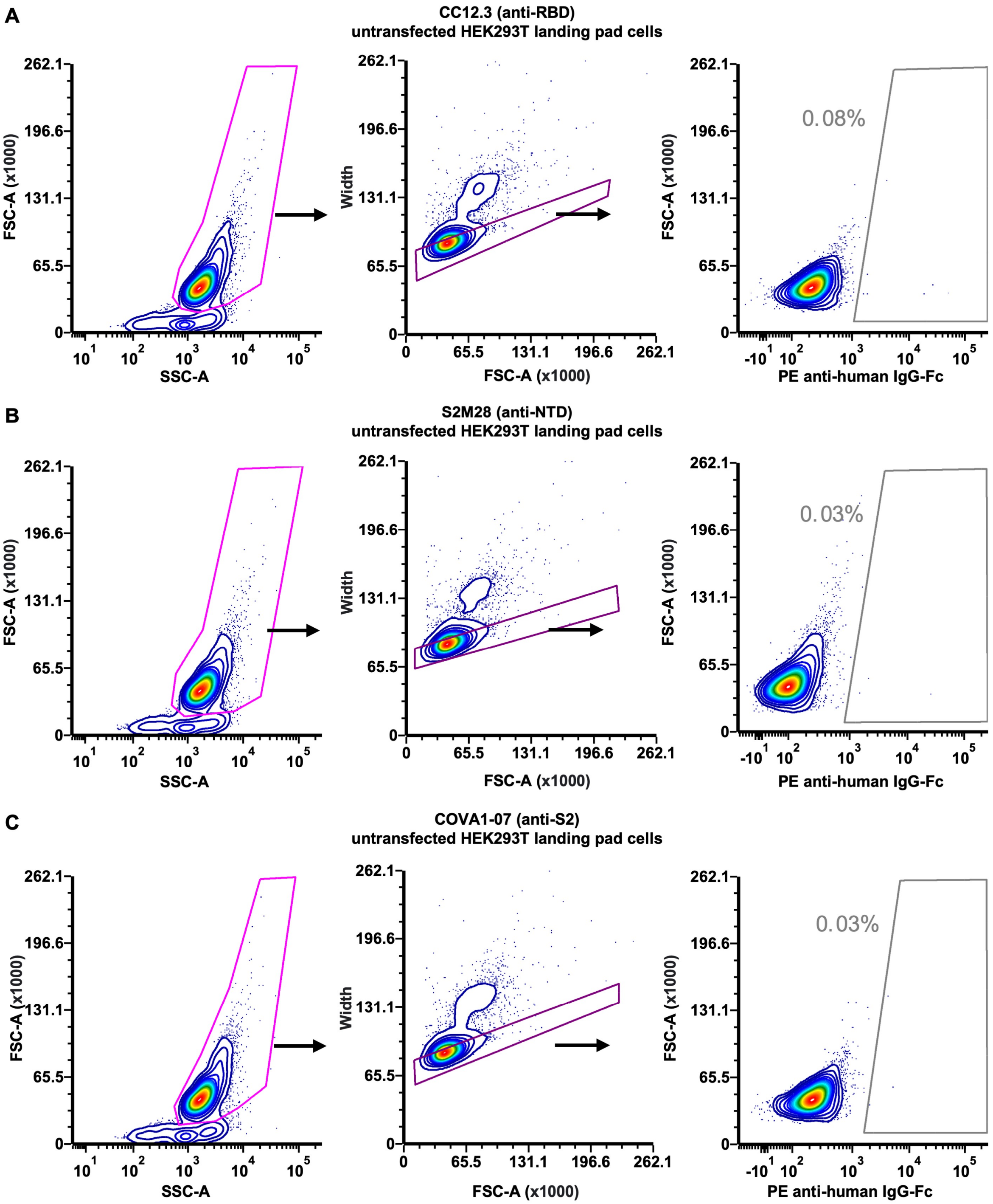


**Supplementary Figure 6. Flow cytometry gating strategy for antibody binding experiment.**

Gating strategy for measuring the binding of S-expressing cells to antibodies **(A)** CC12.3 **(B)**

S2M28, and **(C)** COVA1-07 was setup based on untransfected HEK293T landing pad cells, which serve as a negative control.

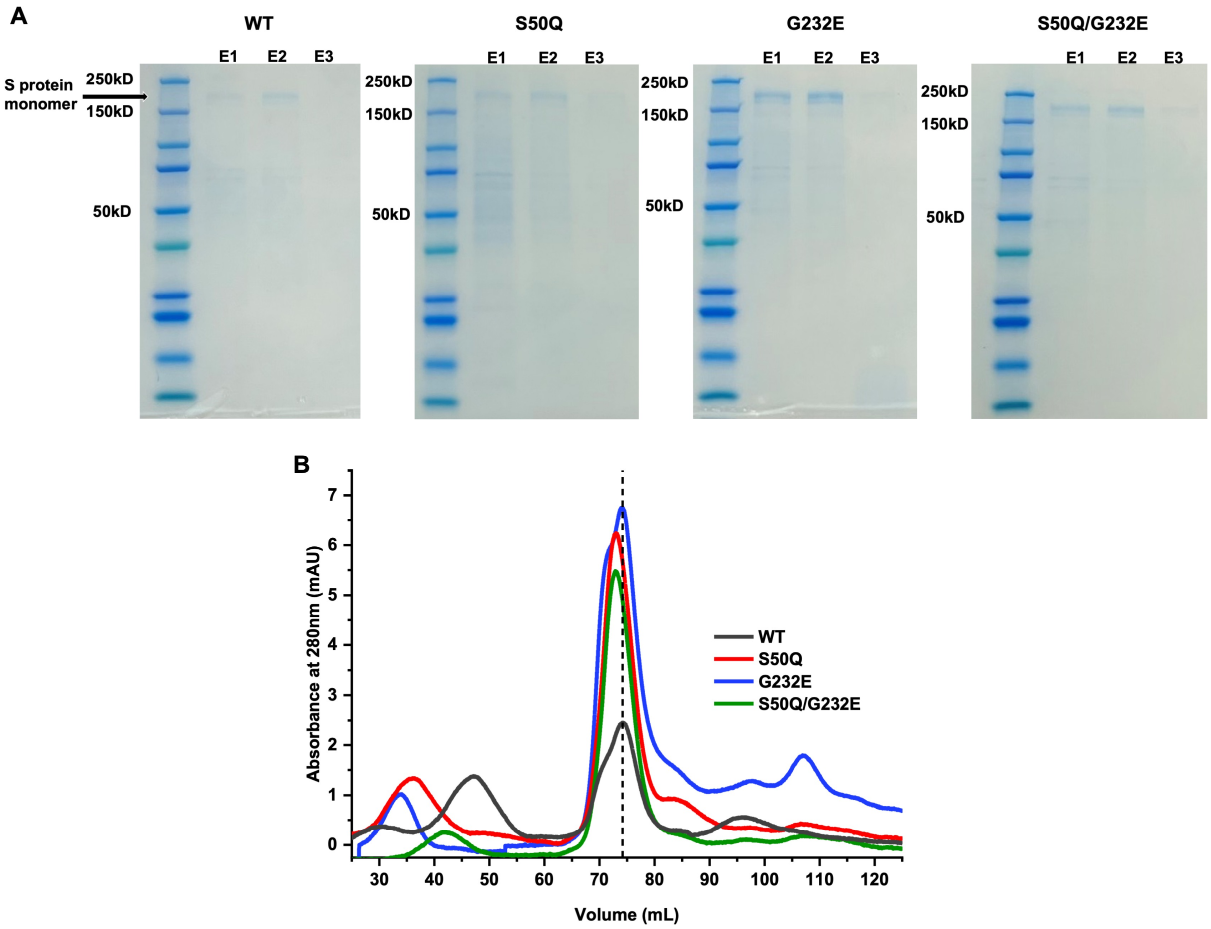


**Supplementary Figure 7. Purification and quality control of recombinantly expressed soluble S proteins. (A)** The SDS-PAGE gel images of affinity purified WT and mutant soluble S proteins are shown. **(B)** The size exclusion chromatographs of the affinity purified WT and mutant soluble S proteins are compared. The dotted line indicates the retention volume of the WT (74.2 mL).

**Table S1. Forward primers for the NTD mutant library construction.**

**Table S2. Reverse primers for the NTD mutant library construction.**

**Table S3. Other primers in this study.**

**Table S4. Experimental data of the expression and fusion assays.**

**Table S5. Raw data of the thermostability assay.**

**Table S6. List of the sarbecovirus strains in the sequence conservation analysis**
